# OsciDrop: A Versatile On-demand Droplet Generator

**DOI:** 10.1101/2021.06.14.448273

**Authors:** Shun Ye, Xu Zheng, Caiming Li, Weihang Huang, Yi Tao, Yanghuan Yu, Limin Yang, Ying Lan, Liang Ma, Shengtai Bian, Wenbin Du

**Author notes:** **Corresponding Author**, **Wenbin Du** – *State Key Laboratory of Microbial Resources, Institute of Microbiology, Chinese Academy of Sciences, Beijing 100101, China; Savaid Medical School, University of the Chinese Academy of Sciences, Beijing 100049, China; College of Life Sciences, University of the Chinese Academy of Sciences, Beijing 100049, China*;, **Xu Zheng** – *State Key Laboratory of Nonlinear Mechanics, Institute of Mechanics, Chinese Academy of Sciences, Beijing 100190, China*;, **Shengtai Bian** – *School of Sport Science, Beijing Sport University, Beijing 100084, China.

## Abstract

Droplet microfluidics is a powerful tool in many biological and clinical applications. Microfluidic chips, such as flow-focusing droplet generators, have been extensively used to high-throughput encapsulate reactions with single-cell and single-molecular resolutions. However, microfabrication is expensive and precision-demanding, preventing it from widespread use in biomedical laboratories and clinical facilities. Herein, we present a versatile chip-free droplet generator, OsciDrop, for generating size-tunable droplets on demand, with high uniformity. OsciDrop segments the fluid flowing out of the orifice of a micropipette tip into droplets by oscillating the tip under the surface of a continuous oil phase. We investigated the factors influencing droplet generation by examining several control parameters. Results show that flow rate, oscillating amplitude, and frequency are key parameters to generate monodisperse droplets on demand. And OsciDrop is able to generate droplets in a flexible and repeatable manner. Importantly, using an optimal asymmetrical oscillation waveform, OsciDrop can controllably generate monodisperse droplets spanning a wide volume range (200 pL - 2 μL). To demonstrate the ability of OsciDrop for chip-free droplet assays, a digital loop-mediated isothermal amplification (dLAMP) was performed to absolutely quantify African swine fever virus (ASFV). The OsciDrop method opens up a feasible and versatile avenue to perform droplet-based assays, exhibiting full accessibility for chip-free droplet microfluidics.

## INTRODUCTION

Droplet microfluidics is an emerging technique in many essential research areas such as biology, medicine, and chemistry due to inherent advantages such as high throughput, minimal sample consumption, low contamination risk, and ease of microenvironment manipulation.^1–5^ By using droplets as microreactors, we can design integrated microfluidic devices to simplify and scale up conventional bench-top experiments performed in flasks, tubes, multiple-well plates, and so on.^6^ Currently, microfabricated devices are primary tools for generating droplets that encapsulate a picoliter to nanoliter volume of reagents and samples.^7–10^ T-junction and flow-focusing designs are commonly used for droplet generation due to geometrical simplicity.^11–16^ Unfortunately, chip-based droplet microfluidics suffers from the following drawbacks: 1) the devices are usually fabricated using time-consuming, labor-intensive photolithography and etching process with cost-prohibitive specialized equipment in cleanrooms;^17–20^ 2) the chip fabrication and operation are complicated, which is a technical barrier for inexperienced researchers;^18,19,21^ 3) it is difficult to precisely define droplet volumes by using multiple pumps and synchronizing several factors (*e.g*., flow rate, fluid viscosity, channel geometry, and surface tension);^20, 22^ These limitations hinder the prevalence of chip-based droplet generation.

Recently, chip-free droplet generation methods, such as centrifugal emulsification^23, 24^ and capillary/needle-based approaches,^18, 20, 22, 23^ emerge as promising alternatives for generating droplets on demand without microfabrication. For instance, using a bench-top centrifuge with the aid of a micro-nozzle array to generate water-in-oil (w/o) droplets has been utilized for digital polymerase chain reaction (PCR), reducing the complexity of droplet assays.^23, 24^ Alternatively, capillary/needle-based methods have been used to produce droplet arrays for *in situ* detection or building biomimetic materials.^24, 25^ We reported interfacial emulsification,^20^ a simple method leveraging a capillary vibrating across the air/continuous phase interface for producing droplets with controllable volumes, and applied this method to synthesize polymeric microparticles,^26^ conduct digital antimicrobial susceptibility testing (dAST),^27^ and perform digital loop-mediated isothermal amplification (dLAMP) for absolute quantification of viruses^24^ and pathogens.^25^ We also demonstrate that oscillating a capillary under the oil surface could segment the aqueous streams flowing out the capillary’s orifice into droplets without the need for localizing air/oil interface precisely.^26^ Similar approaches, such as generating droplets on demand by spinning,^19^ revolving,^16^ or beveled^23^ capillaries/needles, have displayed potential for widespread adoption due to reduced cost and experimental difficulties. Nevertheless, these methods require manual pre-processing of the dispensing capillary (*e.g*., cutting, beveling, heating, and stretching)^18, 20, 23, 27^ or connecting the capillary with tubing using capillary wax or epoxy glue,^18,20^ which is not readily manufacturable. Therefore, a chip-free droplet generation method using easily accessible instruments without manual pretreatment and assembling is attractive. A reliable, on-demand droplet generator practicable for many essential applications (*e.g*., molecular diagnosis,^21^ single-cell analysis,^27^ bacteria/cell cultivation,^28, 29^ nanomaterial synthesis,^30^ etc.) is urgently desired in order to circumvent the drawbacks of current techniques and lower the technical barrier.

Here, we report a versatile chip-free droplet generator, termed as OsciDrop (Fig. 1A), for generating monodisperse droplets on demand, with high uniformity. OsciDrop segments the aqueous fluid flowing out of the orifice of mass-manufactured micropipette tips into droplets by asymmetrically oscillating the tip under the air/oil interface in oil-filled microwells. The generated planar monolayer droplet arrays (PMDAs) on the flat bottom of the microwell facilitate *in situ* reaction and detection. In this work, we developed a mechanical model to analyze the droplet segmentation mechanism and quantitatively delimit different working ranges of the OsciDrop system. We next unveiled primary parameters that influence droplet generation, including flow rate, oscillating amplitude, and frequency. We examined the feasibility of deterministic droplet generation with precise size control spanning picoliter to microliter range. As a proof-of-concept, a dLAMP assay for absolute quantification of African swine fever virus (ASFV) was performed to show OsciDrop’s applicability in molecular diagnostics.

**Figure 1.**
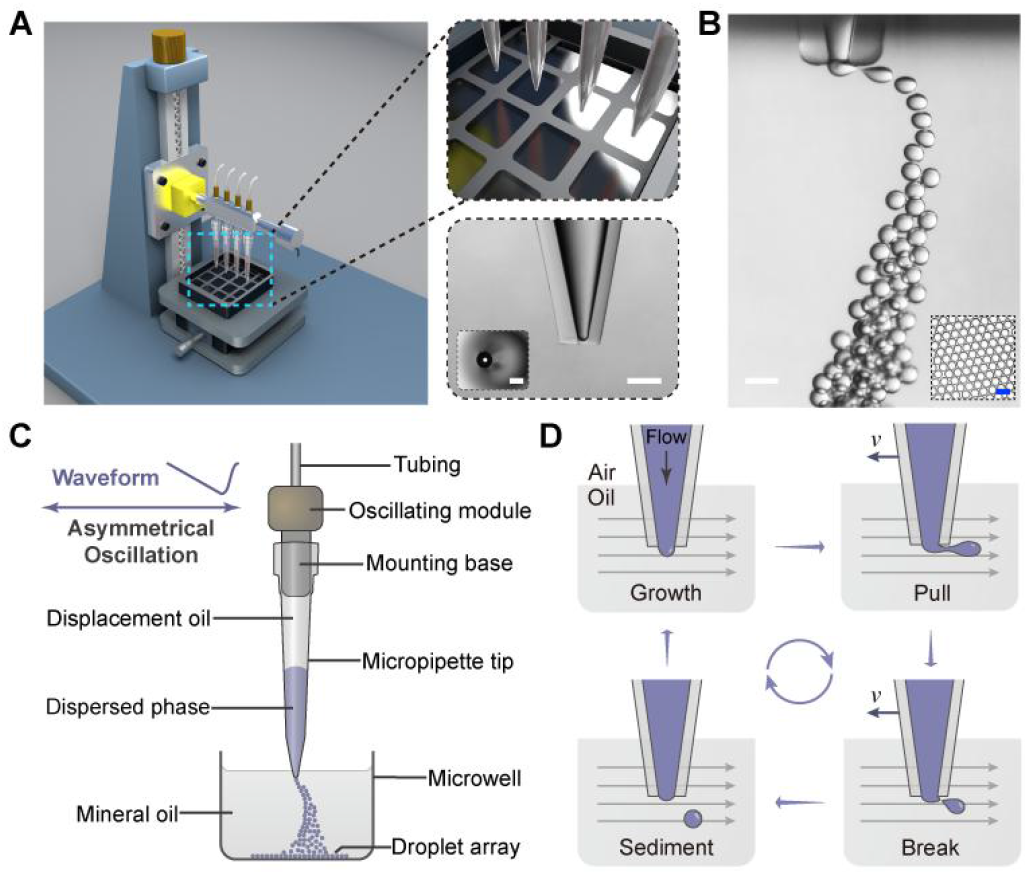
Droplet generation with OsciDrop. (A) Schematic illustrations of the versatile droplet generator consisting of several micropipette tips mounted on the oscillating module, a multi-well plate with oil-filled microwells, and a vertical stage fixed on an optical breadboard. The expended panel shows a magnified view of the micropipette tips and the multi-well plate. Insets are the micrographs of the micropipette tip. Scale bar is 400 μm. (B) Microscopic images of generating water-in-oil (w/o) droplets in a rectangular glass cell and the planar monolayer droplet array (PMDA) at the flat bottom of the microwell (shown in inset). Scale bar is 200 μm. (C) A schematic of an oscillating micropipette tip with its orifice adjusted under the air/oil interface to segment aqueous streams into monodisperse droplets. The droplets sink and form a PMDA subsequently. (D) The schematic shows the working principle and the droplet generation cycle by OsciDrop. The grey arrows indicate the direction of relative motion of the oil phase.

## EXPERIMENTAL SECTION

### Setup of the OsciDrop system

The nanotips (custom-designed micropipette tips with orifice i.d. of ~120 μm), multi-well plates, oscillating module, displacement oil, and droplet generation oil are provided by Dawei Biotech (Beijing, China). Fig. 1A shows the schematic illustrations of the OsciDrop system, which consists of an array of nanotips that attached to an oscillating module, a muti-well plate, and a high-precision vertical stage fixed on an optical breadboard (Zolix, Beijing, China). A waveform generator (RIGOL Technologies, Suzhou, China) was connected with a signal amplifier for controlling the oscillating amplitude and frequency of the oscillating module. Syringe pumps are connected with the oscillating module to infuse displacement oil. The dispersed aqueous phase (TE buffer or LAMP mix) was aspirated and infused through the nanotips at predefined flow rates according to the oscillating frequency and the expected droplet volume and number. The multi-well plate was filled with droplet generation oil.

### Droplet generation by OsciDrop

The OsciDrop system was used to generate monodisperse droplets, as shown in Fig. 1B-C. The nanotip was set to ~ 0.3 mm under the air/oil interface. By dispensing aqueous solution preloaded in the nanotip while the tip was asymmetrically oscillating, the aqueous phase flowing out was segmented into monodisperse droplets (Fig. 1D). We generated droplets with OsciDrop by adjusting several parameters, such as flow rate *Q*, oscillating amplitude *A*, and oscillating frequency *f*.

### Theoretical analysis of droplet segmentation by OsciDrop

To understand the droplet segmentation mechanism of the OsciDrop system, we establish a theoretical model based on the force balance of the droplet. Before the droplet segmentation, the growing droplet will lag behind the oscillating tip due to the viscous drag of the oil phase and form a “neck” of the aqueous stream (Fig. 2A). The involved forces exerting on the droplet include the interfacial tension *F*_σ_, the viscous drag force *F*_v_, the inertial force *F*_i_, the gravity/buoyancy, the kinetic force, and the lift force. After theoretical calculations, only two major horizontal forces *F*_i_ and *F*_v_ are considered to balance the *F*_σ_, which resists the neck break-off; other forces are neglected to simplify the theoretical modeling. Based on the force balance *F*_i_ + *F*_v_ = *F*_σ_, we deduce the equation and determine the working mechanisms (*We*-dominated or *Ca*-dominated) by calculating the *We* number and *Ca* number to comparing the effects of inertial force with viscous force.

**Figure 2.**
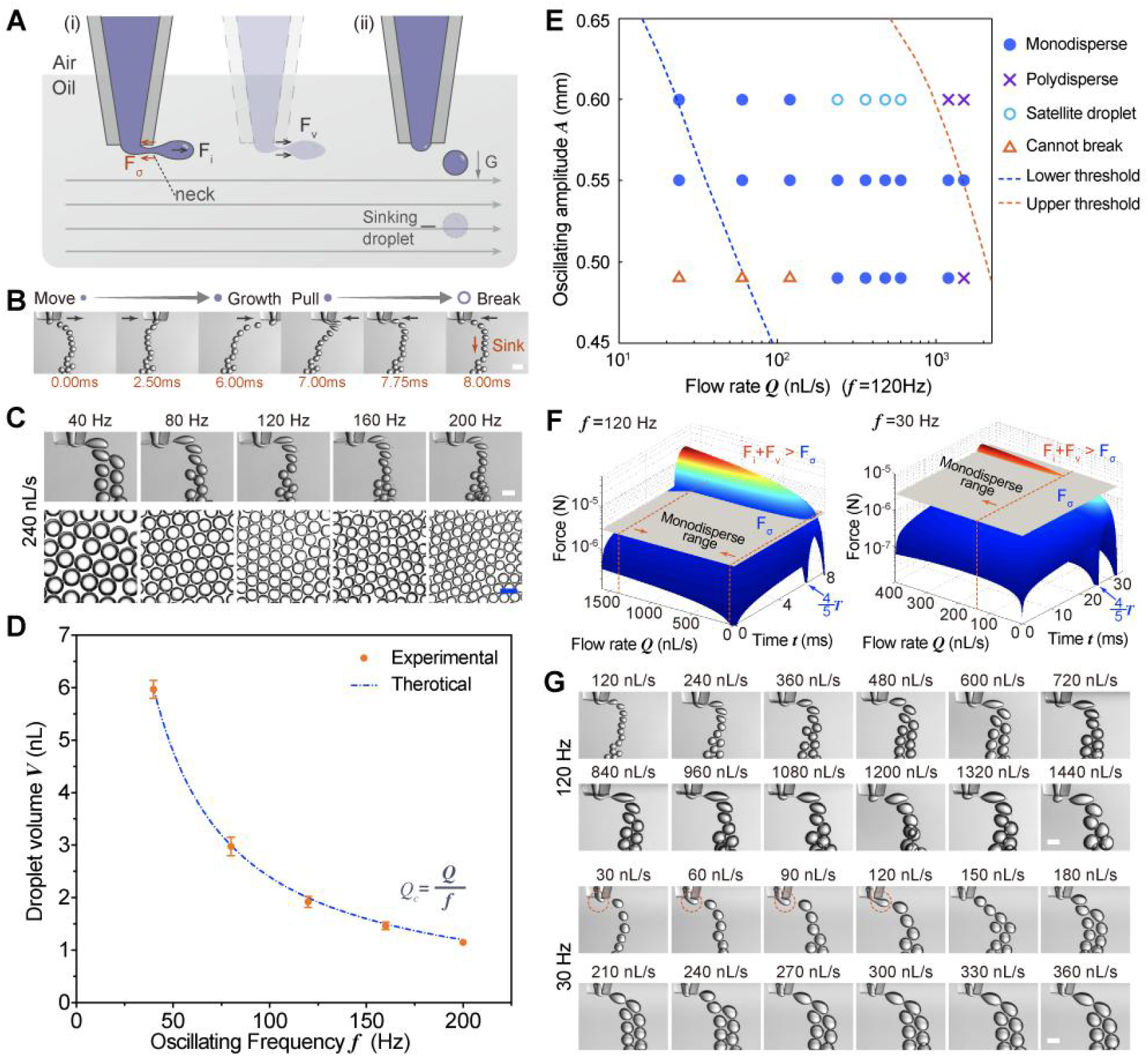
Principle and contributing factors of droplet generation by OsciDrop. (A) Theoretical force analysis of droplet segmentation under the air/oil interface. (i) The force equilibrium among interfacial tension *F*_σ_, inertial force *F*_i_, and viscous drag force *F*_v_ at the “neck” of the aqueous stream. (ii) The droplet sinking after the liquid “neck” break-off. Grey arrows indicate the direction of relative motion of the oil phase. (B) Time-series microscopic images of a representative droplet generation cycle by OsciDrop, including an initial stage (the head of aqueous stream bulges and elongates in the oil phase) and a segmenting stage (the head of aqueous stream is segmented into droplets). Black arrows indicate the moving direction of the micropipette tip under the oil surface. (C) Images of droplet segmentation by OsciDrop under increasing oscillating frequencies *f* from 40 to 200 Hz (flow rate *Q* = 240 nL/s) and the corresponding PMDAs at the flat bottom of microwells. (D) The correlation between the droplet volume *V* and oscillating frequency *f* when *Q* = 240 nL/s. Blue dashed line represents theoretical calculations. (E) Working range map for experimental results (symbols) and theoretical calculations (dashed lines) for showing the relations among *f*, *Q*, and the feasibility of droplet segmentation during one oscillating cycle (*f* = 120 Hz). Blue dots represent desired droplet segmentation. See the legend and Text S6 for details. (F) Theoretical force comparison 3D maps showing the temporal variations of *F*_i_ + *F*_v_ (colorful curved surface) and *F*_σ_ (grey surface) during one period *T* (*f* = 120 Hz, 30 Hz). When the summation of *F*_i_ and *F*_v_ exceeds the *F*_σ_ during segmenting stage (4/5 *T* to *T*), a successful droplet segmentation could be achieved under the corresponding *Q* range. (G) Images of droplet segmentation under increasing *Q*_c_ (*Q*/*f*) (from 1 to 12) (*f* =120 Hz, 30 Hz). The orange dashed circles indicate droplet segmentation needs multiple oscillating circles at small *Q*_c_ values. Scale bar is 200 μm.

To precisely control the droplet segmentation moment by OsciDrop, we analyze the effects of symmetrical and asymmetrical oscillation waveforms on the temporal variations of the inertial force. Afterwards, we plot three-dimensional (3D) force comparison maps for visualizing the temporal variations of breaking forces *F*_i_ + *F*_v_ and interfacial tension *F*_σ_. When the summation of *F*_i_ and *F*_v_ exceeds the *F*_σ_, a droplet segmentation could be achieved. Based on the 3D force comparison, we also define three working ranges for the OsciDrop system, including (1) below the lower threshold flow rate *Q_th-L_*, droplet segmentation cannot complete within an oscillation period as interfacial tension is dominant; (2) between *Q_th-L_* and the upper threshold *Q_th-U_*, the breaking force overcomes interfacial tension in a narrow region during one oscillation period, forming the monodisperse range; (3) beyond *Q_th-U_*, the working range turns to polydisperse as the breaking force and the interfacial tension become comparable in a wide region during one oscillation period. We may obtain chaotic droplet segmentation or satellite droplets in this range. Finally, we plot a phase diagram based on the three working ranges to conclude the proper working condition of OsciDrop (See Text S1 for a detailed version).

### dLAMP

The dLAMP assay for African swine fever virus (ASFV) was custom-designed. EvaGreen (Macklin, Shanghai, China) was used as the fluorescence indicator. A set of four primers was designed using Primer Explorer 4 (https://primerexplorer.jp) (Table S1) and synthesized by InnoGen Biotech (Tianjin, China). The ASFV DNA stock solution was serially diluted with deionized water. The ASFV DNA dilutions (4 μL) were added to the LAMP mix to a final reaction volume of 25 μL. Afterward, a multi-well plate (four rows, each row has four microwells) was used to perform the dLAMP assay, and each reaction was converted into ~18400 1-nL droplets. 25-μL reaction mixtures with serial dilution of ASFV DNA (1x, 4x, 30x, and 100x) were primed into four in-parallel nanotips via pump aspiration, and the OsciDrop system simultaneously produced droplets every four microwells of the multi-well plate, resulting in four repeats for each reaction. In detail, the reaction mix was pushed out of the nanotip at a flow rate of 120 nL/s while the nanotip was oscillating under the air/oil interface at an amplitude of 0.55 mm under 120 Hz asymmetrical oscillation. The oscillating tips generated 4600 droplets in each microwell, which sank to the flat bottom of the microwell of the multiwell plate for producing PMDAs. After sealing the plate with a transparent plate cover, dLAMP assay was performed at 66 °C on a heater for 60 min.

### Data acquisition and statistical analysis

The micropipette tips were imaged using an SMZ 800 stereomicroscope (Nikon, Japan) with a Spot camera (Diagnostic Instruments, USA). A rectangular glass cell filled with droplet generation oil was used to conduct high-speed microscopic imaging during droplet generation with a horizontal objective lens (Venus Optics, Hefei, China). The droplet generation process was recorded via an Optronis CP70 camera (Stemmer Imaging, Germany) controlled by AcutEye V 4.0 software (RockeTECH technology, China). The droplet array images were acquired using a Ti-E inverted fluorescence microscope (Nikon, Japan) equipped with a CoolSNAP HQ^2^ camera (Photometrics, Tucson, USA). The numbers and volumes of droplets were quantified by analyzing the images (200 pL to 100 nL droplets) or the time-series droplet generation images (500 nL to 2 μL droplets) using NIS-Elements software (Nikon, Japan), ImageJ software (NIH, USA), and custom-written Python software. dLAMP experimental results (virus concentrations) were calculated based on Poisson distribution.^27^ Statistical analysis was performed using the GraphPad Prism software (GraphPad Software, USA) and Origin Lab Pro software (OriginLab Corp., USA).

## RESULTS

### Origin and verification of OsciDrop concept

It is technically challenging to generate droplets on demand spanning a broad size range due to geometrical restrictions. This enlightens us to develop a chip-free droplet generator, OsciDrop, which consists of a moving nanotip under a stationary oil phase for segmenting the dispersed aqueous phase to generate droplets. Besides, we designed mass-producible micropipette tips to replace traditional microfluidic chips. To the best of our knowledge, none of the existing techniques leverages micropipette tips to generate monodisperse droplets with controllable volumes since the tip with a large orifice for transferring reagents faces challenges of generating nanoliter droplets. To tackle this challenge, we designed an oscillating module for controllably oscillating the micropipette tip with precise frequencies and waveforms. During the continuous horizontal oscillation, the aqueous phase flowing out of the tip’s orifice is segmented into uniform-sized w/o droplets in sequence at high frequency. Moreover, we hypothesized and validated that the asymmetrical oscillation can outperform the symmetrical oscillation, leading to higher droplet uniformity in a larger dynamic range.

To validate the OsciDrop concept’s feasibility, we established a theoretical model based on the force balance described in *Experimental* and Text S3. In brief, our model unveiled the *We*-dominated droplet segmentation mechanism of the OsciDrop system and establishes the force balance in the horizontal direction (Fig. 2A):

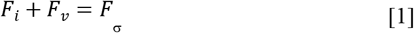

The forces on the left-hand side of Eq. 1 try to detach the droplet from the aqueous stream, while the interfacial tension on the right-hand side resists the neck break-off. When the summation of *F*_i_ and *F*_v_ exceeds *F*_σ_, a successful droplet segmentation will be finished.

Based on our theoretical evaluation, we set up the OsciDrop system and tested the droplet generation using various oscillation waveforms (*e.g*., sine, square, and triangle waves) (see Text S2 and Fig. S1 for details). Results showed that it is technically challenging to generate monodisperse droplets in a controllable fashion using symmetrical oscillations because the summation of *F*_i_ and *F*_v_ may exceed *F*_σ_ twice times during each oscillation cycle (Text S7). Then, we designed an asymmetrical waveform consisting of a long initial stage and a short segmenting stage, which would generate monodisperse droplets. Thus, we design an asymmetrical oscillation waveform, including a long triangle wave (which lasts the first 4*T*/5 in each period) and a short cosinusoidal wave (which lasts the remaining *T*/5 in each period). At an amplitude *A* of 0.55 mm 120 Hz oscillation with a flow rate of 120 nL/s, we successfully produced uniform-sized w/o droplets with OsciDrop. As shown in Fig. 2B, the aqueous stream bulges and elongates during the long initial stage, and the head of the stream is segmented into a droplet during the short segmenting stage, verifying our assumption and the practicability of the OsciDrop concept.

### Mechanism and control parameters of droplet generation

To quantitatively describe the droplet generation process, we define the flow rate within a single oscillating circle as *Q*_c_, which is specified as:

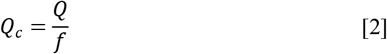

where *Q*, *f* are the flow rate (nL/s) and oscillating frequency (s^−1^), respectively. Under suitable conditions, the droplet generation frequency could be the same as the oscillating frequency, resulting in a produced droplet volume *V* equal to *Q*_c_. To validate the effectiveness of the asymmetrical oscillation, we generated droplets at a fixed flow rate of 240 nL/s with OsciDrop using increasing oscillating frequencies (*i.e*., 40, 80, 120, 160, and 200 Hz; Fig. 2C). We measured the generated droplet volumes by analyzing the images of PMDAs and plotted the results in Fig. 2D. According to Eq. 2, higher oscillating frequencies generated smaller droplets with a fixed *Q* due to decreased *Q*_c_ values. We found that the experimental results of droplet volumes matched well with the theoretical calculations. Thus, it is easy to predict and control generated droplets with precise volumes by OsciDrop under such tested conditions.

To improve the controllability of OsciDrop and dig out the primary control parameters, we further analyze the force balance mentioned above. We deduce the final equation:

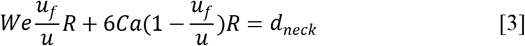

to describe the relation among the *We* number, *Ca* number, aqueous phase injection speed *u_f_*, the oscillating speed *u*, and the generated droplet radius *R* (Text S3). Our calculations have shown that the *We* number is almost two orders of magnitude larger than the *Ca* number, indicating a *We*-dominated droplet segmentation mechanism. In other words, inertial force *F*_i_ is the dominant force determining the droplet segmentation. For the asymmetrical oscillation combining triangle wave and cosinusoidal wave, we deduce the temporal variations of *F*_i_ (see Text S5 for details):

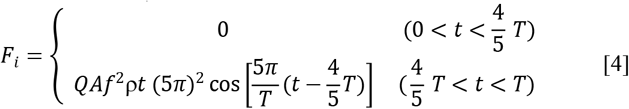

where *t* is the elapsed time. According to Eq. 4, during the initial stage, without the *F*_i_, the *F*_v_ could hardly exceed the *F*_σ_, the head of the aqueous stream bulges and elongates in the oil phase; during the segmenting stage, the droplet detaches due to the dramatically increased *F*_i_, and *F*_i_ is positively proportional to *Q*, *A*, and *f*^2^.

Based on Eq. 4, we suppose that under a fixed *f*, small amplitude *A* means decreased *F*_i_ values during the segmenting stage, resulting in less effective droplet segmenting capacity for generating tiny droplets due to small *Q*·*A* values. In contrast, a too-large amplitude *A* leads to larger *F*_i_ values, resulting in either satellite droplets or poor monodispersity at high flow rates. Thus, a suitable oscillating amplitude *A* is crucial for droplet monodispersity. We tested the droplet generation with OsciDrop by varying flow rate *Q* (24 nL/s to 1320 nL/s) and oscillation amplitude *A* (*i.e*., ~0.5, 0.55, and 0.60 mm), and *f* was set as 120 Hz. Experimental results (symbols) and theoretical prediction (dashed lines) (Text S6) were shown in Fig. 2E: the orange triangles represent the aqueous stream cannot be segmented into droplets during one oscillating circle at low flow rates; the cyan open circles indicate satellite droplet generation; the purple checkmarks represent high polydispersity. We found that 0.55 mm is the most suitable *A* under such tested conditions, which can generate monodisperse droplets (shown as blue dots in Fig. 2E) spanning a wide range of flow rates. We produced droplets from 200 pL to 12 nL by simply changing the flow rate with an amplitude of 0.55 mm under 120 Hz oscillation (will be described later).

In addition to the oscillating amplitude *A*, the oscillating frequency *f* also plays a crucial role in determining the droplet segmentation. Theoretically, the reduced *f* leads to dramatically decreased *F*_i_, resulting in limited ability to generate tiny droplets. To predict the droplet segmentation and delimit different working ranges, we calculated the temporal variations of *F*_i_ + *F*_v_ and *F*_σ_ (Text S1 and S6). The force variations of 120 Hz and 30 Hz asymmetrical oscillations during one period *T* are compared in Fig. 2F. According to our theoretical prediction, the summation of *F*_i_ and *F*_v_ (shown as colorful curved surface) dramatically reduces in response to decreased frequency, especially during the segmenting stage, resulting in ineffective droplet segmentation under small *Q*_c_ values. To verify our theoretical evaluation based on Eq. 3 and 4, we generated droplets using increasing *Q*_c_ (1 to 12) with an amplitude *A* = 0.55 mm under 120 Hz and 30 Hz oscillations, respectively (Fig. 2G). Results show that it is facile to generate monodisperse droplets with increasing *Q*_c_ under 120 Hz oscillation, whereas it is unable to produce tiny droplets when *Q*_c_ ≤ 4 under 30 Hz oscillation. The monodisperse ranges under two different oscillating frequencies matched very well with the theoretical prediction, verifying the essential role of oscillating frequency *f* in droplet generation by OsciDrop.

### Predictable on-demand droplet generation using OsciDrop

After understanding the working principle of OsciDrop and seeking out three control parameters, we then examined the predictability and flexibility of OsciDrop for generating monodisperse droplets. Keeping the oscillating amplitude *A* constant at 0.55 mm under 120 Hz oscillation as described above, we observed that increasing flow rate *Q* from 24 nL/s to 1440 nL/s enlarged the droplet size predictably, as shown in Fig. 3A. Then, we also measured the generated droplet volume *V* and plotted the results in Fig. 3B. The linear correlation between the droplet volume *V* and flow rate *Q* (R^2^=0.9976) was consistent with the theoretical calculation by Eq. 2. Thus, it is easy to obtain expected droplet volumes (200 pL to 12 nL) by simply adjusting the flow rates within this monodisperse range, reflecting high predictability and controllability of the OsciDrop method.

**Figure 3.**
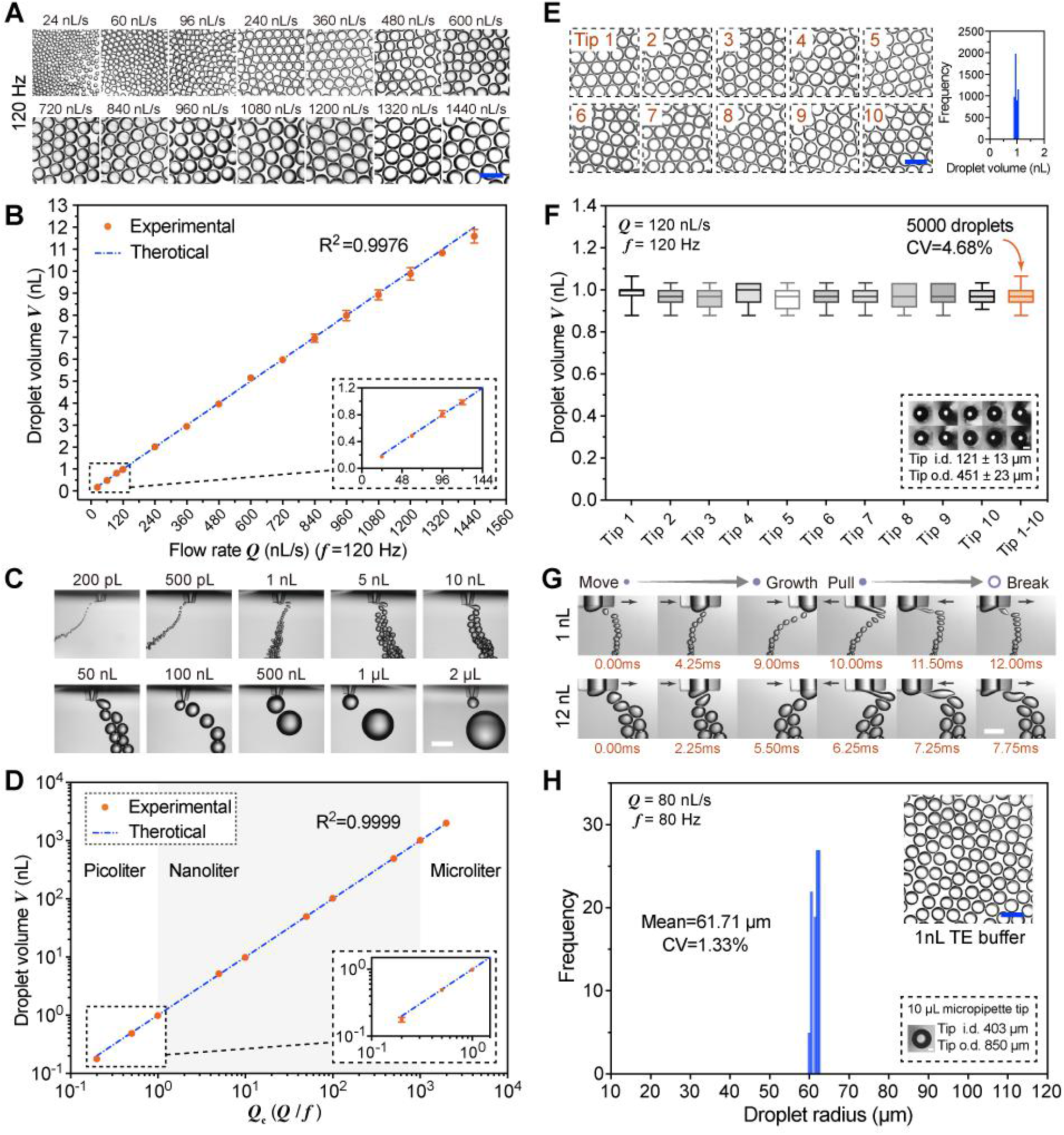
Droplet generation by OsciDrop is predictable, controllable, flexible, repeatable and robust. (A) At an amplitude *A* of 0.55 mm under 120 Hz oscillation, droplet size predictably enlarges with increasing flow rate *Q* (24 nL/s to 1440 nL/s). Scale bar is 400 μm. (B) The linear correlation between the droplet volume *V* and flow rate *Q* at a fixed *f* of 120 Hz. Blue dashed line represents theoretical calculations. (C) Size-tunable generating monodisperse pico/nano/microliter droplets using OsciDrop by simultaneously adjusting three control parameters (*i.e., Q, A, f*). Scale bar is 800 μm. (D) The linear correlation between the generated droplet volume *V* and calculated *Q*_c_ value. (E) PMDAs generated at a flow rate *Q* of 120 nL/s and a frequency *f* of 120 Hz by using ten micropipette tips with different inner/outer diameter (i.d./o.d.). Produced 5,000 droplets’ volume distribution showing that they are the same size with droplet volume *V* of approximately 1 nL. (F) The volume distribution of droplets generated using Tip 1 to Tip 10 (shown in inset), and 5,000 droplets’ volume coefficient of variation (CV) is 4.68%. (G) Time-series microscopic images of the droplet generation cycle (expected volumes of 1 nL and 12 nL) with OsciDrop using a standard 10 μL micropipette tip. Black arrows indicate the moving direction of the micropipette tips. Scale bar is 400 μm (H) The radius distribution of droplets generated at a flow rate *Q* of 80 nL/s and a frequency *f* of 80 Hz (expected volume of 1 nL (radius of 62.04 μm)) using a standard 10 μL micropipette tip (shown in inset), showing OsciDrop’s robustness. Scale bar is 200 μm (E, F, and H).

Although size-tunable droplet generation can be easily achieved by adjusting flow rates, it is technically challenging to produce monodisperse droplets spanning a wide volume range (*e.g*., from picoliter to microliter range) with fixed oscillating amplitude and frequency. Therefore, we next investigated the flexibility of OsciDrop for generating pico/nano/microliter droplets that conventionally difficult to achieve in either chip-based or chip-free methods. By adjusting three primary control parameters simultaneously, including *Q*, *A*, *f* (Table S2), we successfully generated monodisperse droplets spanning picoliter to microliter range (*i.e*., 200 pL, 500 pL, 1 nL, 5 nL, 10 nL, 50 nL, 100 nL, 500 nL, 1 μL, and 2 μL), as shown in Fig. 3C. It seems to be complicated to control the droplet volume by synchronizing three parameters. In truth, it is simple and straightforward: we calculate the *Q*_c_ values before the experiments according to expected droplet volumes and decrease the oscillating frequency when expected droplet volumes increase to facilitate generating large droplets. Fig. 3D shows that the linear correlation between the generated droplet volumes *V* and calculated *Q*_c_ value matched near-perfectly with theoretical calculations (R^2^=0.9999), validating the flexible droplet generation with tunable volumes spanning 5 orders of magnitude. This unique feature may further extend the range of OsciDrop’s applications in many essential research areas.

### Validation of OsciDrop’s repeatability and robustness

In contrast to standard photolithography and etching process for microchip fabrication with high precision, mass-produced nanotips by conventional plastic injection molding are relatively inaccurate, resulting in uncontrollable variations in the inner/outer diameter (i.d./o.d.) of the tip’s orifice. In other words, repeatable droplet generation by OsciDrop using mass-produced tips with different i.d. and o.d. would be an indispensable prerequisite for widespread use in academia and clinics, as well as successful industrialization. Therefore, we generated droplets (expected volume *V* is 1 nL) using ten randomly picked nanotips. The generated PMDAs were then recorded and quantitatively analyzed (Fig. 3E). At a flow rate of 120 nL/s and oscillating frequency of 120 Hz, 5,000 droplets generated by ten mass-produced tips were of the same size with volumes of approximately 1 nL and a coefficient of variation (CV) of 4.68%. Fig. 3E *right* and 3F plot the volume distribution of droplets produced by Tip 1 to Tip 10 (i.d. 121 ± 13 μm, o.d. 451 ± 23 μm) (tip micrographs were shown in inset), which demonstrates high repeatability and considerable potential in digital molecular diagnostics in which generating droplets with precise size control is required.

Although the above results reflect excellent repeatability of OsciDrop, the customized nanotips having an smaller i.d. are not readily accessible in many laboratories, which limits the usability of OsciDrop due to lack of consumables. Therefore, we examined the robustness of OsciDrop by determining whether or not it could generate monodisperse droplets using standard micropipette tips available in common laboratories. We produced droplets (expected volumes of 1 nL and 12 nL) using a standard 10 μL micropipette tip (i.d. 403 μm, o.d. 850 μm) by simultaneously fine-tuning three control parameters. As shown in Fig. 3G, we successfully generated uniform-sized monodisperse droplets with expected volumes. We analyzed the PMDA generated at a flow rate of 80 nL/s under 80 Hz asymmetrical oscillation using a standard 10 μL tip and plotted the produced droplets’ radii distribution (Fig. 3H). We found that although the diameter of droplets (approximately 124 μm) is far smaller than the inner diameter of the 10 μL tip (approximately 400 μm), the head of the aqueous stream still can be segmented into droplets (see Text S8 and Fig. S3 for details). This pilot experiment unveils the near-limitless potential of our OsciDrop system: in the future, every biology, chemistry laboratories having 10 μL micropipette tips could use OsciDrop to perform chip-free droplet microfluidic experiments for high-throughput and parallel encapsulating biochemical reactions, reducing reagent consumption, as well as breaking through technical barriers for beginner researchers.

### dLAMP ASFV detection using OsciDrop

The recent spread of African swine fever virus (ASFV) has caused animal pandemic in many countries, and resulted in severe economic losses. To allow rapid and high sensitive detection of ASFV, we performed a dLAMP assay for absolute quantification of African swine fever viruses (ASFVs) as schematically illustrated in Fig. 4A. The OsciDrop system utilizes four parallel channels to generate monodisperse droplets that encapsulates LAMP reactions. In brief, we first serially diluted ASFV DNA stock solution and obtained four dilutions of 1x, 4x, 30x, and 100x, followed by 4-channel generating droplets in an oil-filled multi-well plate by simultaneously oscillating four nanotips under the oil. The multi-well plate was then sealed by a plate cover and then incubated at 66 °C for 1 h on a heater for amplification, followed by fluorescence imaging and analysis to calculate the concentration of ASFVs. Fig. 4B shows the end-point fluorescent images of dLAMP reaction solutions with different dilution factors (*i.e*., 1, 4, 30, and 100). Strong fluorescence signals were obtained after amplification for positive droplets and displayed a two-fold change between positive and negative droplets. The measured concentrations were linear with the dilution factor with an R^2^ of 0.9975 (Fig. 4C) and a detection limit of ~3 copies/μL, verifying the reliable performance of OsciDrop in molecular diagnosis with user-friendly operations.

**Figure 4.**
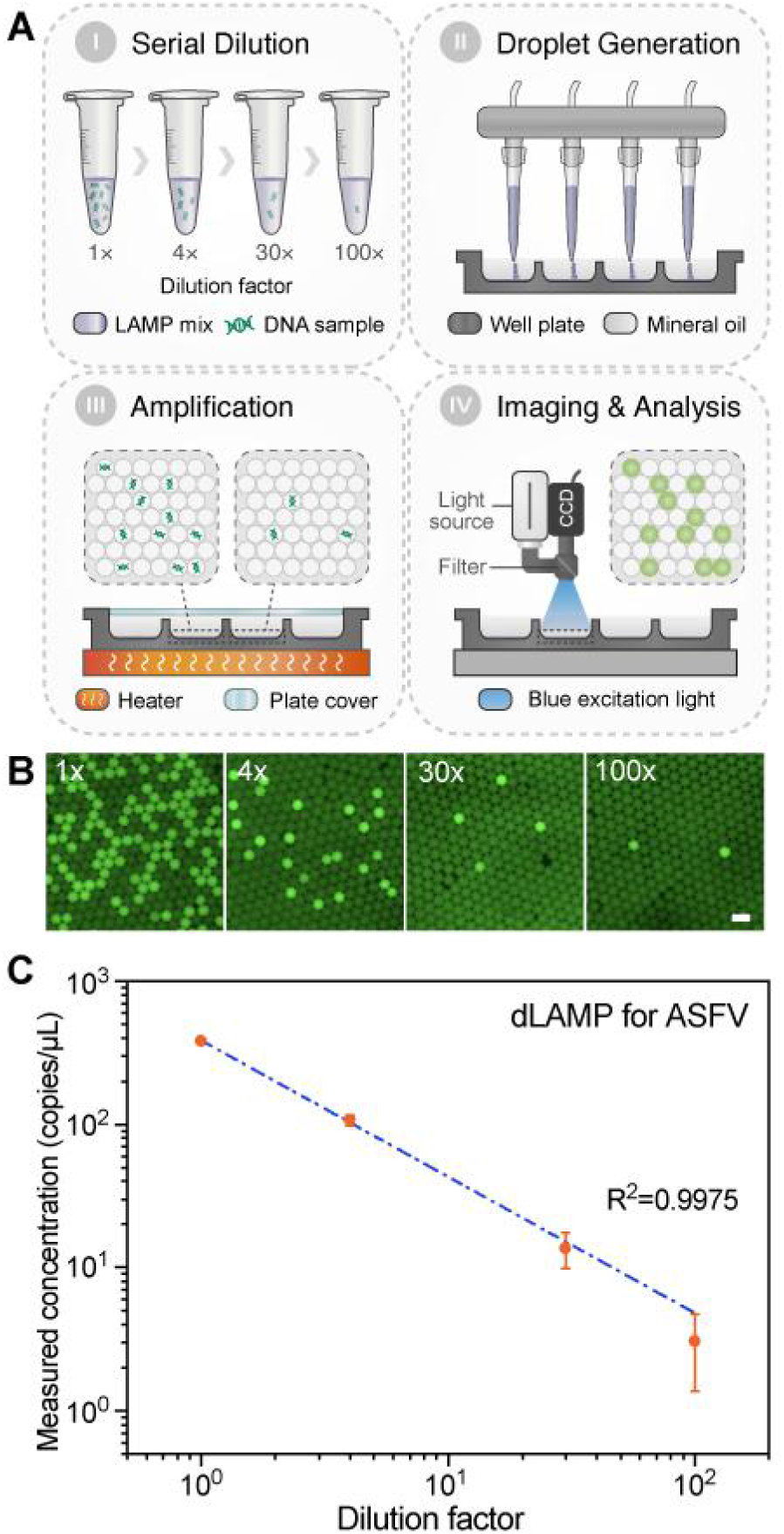
Multichannel droplet generation with OsciDrop for dLAMP. (A) A schematic illustration of the dLAMP workflow, including serial dilution of the African swine fever virus (ASFV) DNA stock solution, in-parallel four-channel droplet generation for sample segmentation, isothermal amplification, and fluorescence detection and quantitative analysis of the concentration of ASFV DNA. (B) End-point fluorescent images of PMDAs in dLAMP assays of serial dilutions (1x, 4x, 30x, and 100x) of the ASFV samples. Scale bar is 200 μm. (C) The linear correlation (blue dashed line) between the measured concentration and the dilution factor of the DNA samples.

## DISCUSSION

Microfluidics is widely utilized in droplet generation and manipulation,^2^ such as reaction encapsulation,^31^ droplet coalescence,^32^ and droplet-based single-cell sorting,^33^ due to its remarkable capacity for performing in-parallel, high-throughput, contamination-free biochemical assays.^3, 4^ Generating monodisperse droplets with precise size control to encapsulate samples and reagents, the foremost and crucial step of sample segmentation, have been achieved using chip- based and chip-free microfluidics.^34, 35^ Take chip-free approaches, for example. Tang *et al*.^19^ reported an off-chip droplet generator enabled by a 34G needle and a spanning conical frustum; however, the large conical frustum limited its use in small containers, such as centrifuge tubes. Although alternative methods, using a revolving needle for generating microdroplets in a centrifuge tube, have been presented,^18, 21^ they induced strong vortices that consequently undermine their utilities in miniaturized containers like microwells of the multi-well plate due to generated droplets could be smashed by vortices.^21^ Other two capillary-based approaches circumvented this limitation but showed narrow monodisperse ranges and difficulties in droplet size control.^20, 36^ The OsciDrop method, which oscillates multiple micropipette tips under the air/oil interface to segment aqueous streams into droplets, is able to generate pico/nano/microliter droplets on demand in multi-well plates Although the OsciDrop system has successfully demonstrated the capability of size-tunable generating droplets with a wide dynamic range (from picoliter to microliter range) under asymmetrical oscillation, we are still attempting to optimize and enhance the performance of OsciDrop in many aspects. We first make an effort to widen the dynamic volume range of droplet generation by OsciDrop. We reduce the *Q*_c_ value to generate 10 pL and 100 pL droplets successfully using a high-precision infusion pump (PHD Ultra, Harvard Apparatus, USA) (Fig. S4). Further, the effect of tapered-tip geometry on the droplet generation using OsciDrop was also investigated. We find that it can facilitate generating picoliter droplets (Text S9 and Fig. S5). In addition to the dynamic volume range, we also attempted to generate droplets using other oscillation waveforms. Fortunately, we successfully generate 1 nL to 12 nL droplets using sine wave (Text S7 and Fig. S6) by simultaneously adjusting three control parameters; however, droplet generation by sinusoidal oscillation is more complex and difficult to control. Further improvement in OsciDrop’s performance could be made by fine-tuning control parameters, optimizing oscillation waveforms, as well as using highly precise pumps.

In recent years, droplet-based microfluidic devices have been widely applied in the absolute quantification of nucleic acids with digital PCR/LAMP techniques,^37, 38^ such as self-digitalization (SD) LAMP chip,^39, 40^ interfacial printing system,^22^ and centrifugal micro-nozzle array.^24^ Besides, many commercial systems emerge as research tools. Based on the OsciDrop concept, we are working on commercializing an all-in-one OS-500 digital PCR instrument, which integrates a robotic automation unit and performs sample loading, droplet generation, *in situ* amplification, multichannel fluorescence detection, and data analysis & visualization in a fully automatic manner, reducing the operational complexity of precision molecular diagnostics.

## CONCLUSION

In this paper, we established OsciDrop, a simple, reliable, flexible, versatile droplet generating method, using mass-fabricated micropipette tips to achieve deterministic droplet segmentation under asymmetrical oscillation. This novel OsciDrop system benefits from the controllable inertial force originated from the asymmetrical oscillation, and thus works in a *We*-dominated range different from previous techniques. The feasibility of size-tunable generating pico/nano/microliter droplets with high uniformity was achieved by adjusting the flow rate, oscillating amplitude, and frequency. Several attractive features of droplet generation by OsciDrop, such as predictability, controllability, repeatability, and robustness, were also verified by wet-lab assays, which may unleash the power of chip-free droplet microfluidics. The unique feature of OsciDrop, especially high flexibility, would be beneficial for addressing a critical challenge, generating monodisperse droplets on demand spanning picoliter to microliter range, in droplet microfluidics. Such a wide dynamic droplet generation range enables numerous applications in biology, chemistry, and medicine.

## ASSOCIATED CONTENT

### Supporting Information

The Supporting Information is available free of charge at: https://doi.org/10.1101/2021.06.14.448273

Detailed version of theoretical analysis of droplet segmentation; examining the feasibility of generating droplets using different oscillation waveforms; deduction of the force balance equation; *We*-dominated mechanism; force analysis for the asymmetrical oscillation; discussion on force plot and three working ranges; performance of sinusoidal oscillation and corresponding force plot; droplet generation using the standard 10-μL micropipette tip; investigation of the effect of wall thickness on the droplet generation using an one-side tapered tip with OsciDrop; LAMP primers used in ASFV detection; Control parameters for generating monodisperse droplets using OsciDrop (PDF)

### Notes

Dr. Du is a co-founder, consultant, and director and has stock ownership from Dawei Biotech., outside the submitted work.

## ACKNOWLEDGMENTS

This work was supported by the National Natural Science Foundation of China (21822408, 12072350, 91951103, and 11832017), the CAS Key Research Program of Frontier Sciences (QYZDB-SSW-JSC036, QYZDB-SSW-SMC008), the CAS Strategic Priority Research Program (XDB22040403), and National Key Research and Development Program of China (2016YFE0205800, 2018YFC0310703).

