## Supporting Information for "OsciDrop: A Versatile On-demand Droplet Generator"

### Supporting Text

#### Text S1. Detailed version of theoretical analysis of droplet segmentation by OsciDrop

To understand the droplet segmentation mechanism of the OsciDrop system, we develop a theoretical model based on the force balance of the droplet. The oscillation of the current OsciDrop system is along horizontal direction. When the aqueous phase is flowing out of the micropipette tip, the growing droplet will be distorted by the horizontal shear flow of the oil phase. During the droplet segmentation, the droplet lags behind the oscillating tip due to the viscous drag of the oil phase, forming a "neck" of the aqueous stream (Fig. 2A). The involved forces exerting on the droplet during droplet segmentation include: the interfacial tension $F_\sigma$, the viscous drag force $F_v$, the inertial force $F_i$ originated from the acceleration/deceleration of the tip oscillation, the gravity/buoyancy of the droplet, the kinetic force owing to the flow rate of the aqueous phase, and the lift force due to the pressure difference of the shear flow of the oil phase. The vertical forces like gravity/buoyance are neglected as the droplet segmentation by OsciDrop occurs in the horizontal direction. The kinetic force is also ignored in our analysis, because it is much smaller than the interfacial tension $F_\sigma$ and the viscous drag force $F_v$. We stress that the viscous drag force from the oil phase and the inertial force originated from the tip oscillation are two major effects causing the droplet segmentation when they exceed the interfacial tension at the neck. Based on the force balance $F_i + F_v = F_\sigma$, we deduce a final equation to describe the relation (Text S3) among the aqueous phase injection speed $u_f$, the oscillation speed $u$, and the generated droplet radius $R$:

$$We\frac{u_f}{u}R + 6Ca(1-\frac{u_f}{u})R = d_{neck} \quad [1]$$

where $We$ and $Ca$ are the Webber number and capillary number respectively, and $d_{neck}$ is the characteristic dimension of the neck region. The $We$ number, $We = \frac{\rho R_{tube}^2 uTa}{\sigma R}$, is the ratio between the inertial force and the interfacial tension, while the capillary number $Ca$, $Ca = \frac{\eta u}{\sigma}$, defines the ratio between the viscous force and the interfacial tension, where $R_{tube}$ is the tube radius in the tip, $\rho$, $\sigma$, and $\eta$ are the fluid density, the tension of the aqueous/oil interface, and the oil viscosity, respectively, and $T$ and $a$ are the period and acceleration of the oscillation. These two dimensionless numbers unveil that the droplet segmentation is determined by $We$-dominated and $Ca$-dominated two ranges. Previous design of spinning emulsion generator works in the $Ca$-dominated range, in which the force balance is simplified as $F_v = F_\sigma$. In contrast, in the current OsciDrop system where accelerated/decelerated oscillation introduces the inertial force $F_i$, the droplet segmentation is dominated by the $We$ number instead (Text S4). In our experiments, the $We$ number (~O(10)) is almost two orders of magnitude larger than the $Ca$ number (~O(0.5)), indicating the significant contribution of the oscillation inertial force $F_i$.

We find that the OsciDrop system can precisely determine the moment when droplet segmentation occurs by adjusting the waveform of the oscillation and varying the inertial force $F_i$. We design an asymmetrical oscillation (Fig. S2) by combining a long triangle wave (called initial stage that lasts the first $4T/5$ in each period) plus a short cosinusoidal wave (called segmenting stage that lasts the remaining $T/5$ in each period). We will show in the *Results* section and Text S4 and S5 that this oscillation is very effective to achieve deterministic droplet segmentation with highly uniform droplet size in every single oscillation period. We calculate the temporal force variation under such asymmetrical oscillation (Fig. 2F, with oscillation frequency $f$ = 120 Hz and 30 Hz), by which we can estimate when the summation of the inertial force $F_i$ and the viscous drag force $F_v$ exceeds the interfacial tension $F_\sigma$. In the Fig. 2F, the interfacial tension $F_\sigma$ is shown by the grey horizontal plane as it is independent of flow rate $Q$ and elapsed time $t$, whereas the colorful curved surface represents the summation of the breaking force $F_i + F_v$. The droplet segmentation occurs in the segmenting stage $t > 4T/5$ when the colorful surface $F_i + F_v$ is higher than the grey plane $F_\sigma$. When the oscillation frequency $f$ = 120 Hz, the breaking force $F_i + F_v$ could be one order of magnitude larger than the interfacial tension, indicating a deterministic effect for droplet segmentation. While if the frequency is reduced to 30 Hz, the breaking force is only slightly larger than $F_\sigma$. We emphasize that there are a lower threshold flow rate $Q_{th-L}$ and an upper threshold flow rate $Q_{th-U}$, as being highlighted by the dashed lines in Fig. 2F, between which we achieve perfectly monodisperse range. Thus, based on the theoretical analysis, we define three working ranges for the OsciDrop system: (1) Below $Q_{th-L}$, droplet segmentation cannot complete in one period as interfacial tension is dominant. (2) Between $Q_{th-L}$ and $Q_{th-U}$, the breaking force overcomes interfacial tension in a narrow region of the segmenting stage, forming the monodisperse range. (3) Beyond $Q_{th-U}$, the working range turns to polydisperse as the breaking force and the interfacial tension become comparable even in the initial stage. We may obtain chaotic droplet segmentation or satellite droplet in the polydisperse range. Based on the three working ranges, we plot a phase diagram to conclude the proper working condition (Fig. 2E). We will show in *Results* that the theoretical phase diagram is in good agreement with the experimental observation (Fig. 2E and Text S6).

**Text S2. Examining the feasibility of generating droplets using different oscillation waveforms**

Based on theoretical assumptions, we examine the feasibility of generating monodisperse droplets using different oscillation waveforms, including sine, square, and triangle waves. Part of results are shown in Fig. S1. We find that it is difficult to generate monodisperse droplets in each period using symmetrical oscillation. Generally, for these symmetrical oscillations, under the small oscillating amplitude or oscillating frequency, the $F_i$ value is relatively small and shows difficulty in overcoming the interfacial tension (see *Experimental* for details). While the oscillating amplitude and frequency are increased, the $F_i$ may exceed the interfacial tension twice during one oscillating period, leading to poor monodispersity and difficulty in controlling the segmenting moment (will be explained in Text S6). We simply explain the shortcomings of these symmetrical oscillations. For instance, for sinusoidal oscillation (see Text S6 for details), the inertial force $F_i$ is usually smaller than that of an asymmetrical oscillation, which causes difficulty to generate uniform droplets with small volumes. If we increase oscillation amplitude or frequency to overcome such problem, we will see nonuniform droplet segmentation happening twice in one oscillation period as the sinusoidal wave has two moment reaching the maximum acceleration $a$ in each period. For square wave, inertial force $F_i$ only exists during the instant switch from positive position to negative one, and *vice versa*. Theoretically, at this moment the acceleration $a$ and the inertial force $F_i$ are both infinite, whereas in practice $a$ and $F_i$ are quite large but dependent on the oscillator's property. This causes chaotic droplet segmentation and undesirable polydispersity. The square wave also leads to nonuniform droplet segmentation happening twice in one oscillation period as it has two instant switch in one period. For the similar reasons, triangle wave is not a suitable choice too. Thus, we propose an asymmetrical oscillation, consisting of a long initial stage and a short segmenting stage, would generate monodisperse droplets in a controllable manner. Results show that our asymmetrical waveform exhibits excellent performance for generating monodisperse droplets (Fig. S1).

**Text. S3. Deduction of the force balance equation**

During droplet segmentation in the OsciDrop system, there are following forces exerting on the droplet: the interfacial tension $F_\sigma$, the viscous drag force $F_v$, the inertial force $F_i$ originated from the acceleration/deceleration of the tip oscillation, the gravity/buoyancy of the droplet, the kinetic force owing to the flow rate of the aqueous phase, and the lift force due to the pressure difference of the shear flow of the oil phase. The droplet segmentation of the OsciDrop system occurs under the horizontal oscillation, so the forces in the vertical direction are neglected. The droplet is considered to be spherical. The three horizontal forces thus establish the force balance:

$$F_i + F_v = F_\sigma \quad \text{[S1]}$$

The forces on the left-hand side of Eq. S1 try to detach the droplet from the aqueous stream, while the interfacial tension on the right-hand side resists the break-off. When the breaking force $F_i$ + $F_v$ exceeds $F_\sigma$, droplet segmentation will complete.
The inertial force can be expressed based on the acceleration $a$ of the oscillation:

$$F_i = ma = \rho QTa = \rho\pi R_{tube}^2 u_f Ta \qquad [S2]$$

where $m$ is the droplet mass, $\rho$ is fluid density, $Q$ is the flow rate of the aqueous phase, $u_f$ is the flow speed of the aqueous phase, $R_{tube}$ is tube radius of the micropipette orifice, and $T$ is the oscillation period. The viscous drag force $F_v$ from the oil phase can be described based on the theory of the Stokes drag force:

$$F_v = 6\pi\eta R(u - u_f) \qquad [S3]$$

where $\eta$ is the oil phase viscosity, $R$ is the droplet radius, and $u$ is the oscillation speed. The interfacial tension is exerted at the neck region, which is written as:

$$F_\sigma = \pi\sigma d_{neck} \qquad [S4]$$

where $\sigma$ is the interfacial tension coefficient, $d_{neck}$ is the diameter of the neck region. Substituting Eqs. S2 - S4 into Eq. S1, we obtain:

$$\rho\pi R_{tube}^2 u_f Ta + 6\pi\eta R(u - u_f) = \pi\sigma d_{neck} \qquad [S5]$$

Both sides of Eq. S5 are divided by $\pi\sigma$, and we deduce the final equation (Eq. 1 in the manuscript) by defining the *We* number $We = \frac{\rho R_{tube}^2 uTa}{\sigma R}$ and the capillary number $Ca = \frac{\eta u}{\sigma}$:

$$We\frac{u_f}{u}R + 6Ca(1 - \frac{u_f}{u})R = d_{neck} \qquad [S6]$$

The *We* number and *Ca* number unveil two different ranges of the droplet segmentation respectively: inertial force-dominated range and viscous force-dominated range.

**Text S4. Novel *We*-dominated mechanism different from previously *Ca*-dominated mechanism**
We stress that the mechanism of our OsciDrop system is totally different from the previous spinning/revolving droplet generator [1-2]. Herein, we will clarify their mechanisms by quantifying the dimensionless *We* and *Ca* numbers based on Eq. S6. For spinning/revolving droplet generator, as the uniform circular motion of the droplet doesn't introduce the inertial force, the *We* number is zero and the droplet segmentation is only determined by the *Ca* number, indicating a viscous force-dominated mechanism. Thus, Eq. S6 can be simplified as:

$$R = \frac{d_{neck}}{6Ca(1 - {}^{u_f}\!/_{u})} \qquad [S7]$$

which is similar to the theoretical result in ref [1]. The reported spinning speed is more than 1000 rpm, corresponding to a *Ca* number in the range of 1 - 5.
However, in the OsciDrop system, the inertial term governed by the *We* number becomes dominated. Based on the definition of the *We* and *Ca* numbers, we calculate their values to compare their effects. In the calculation, the values of all parameters are the same as those used in our experiments, which are listed below. The interfacial tension coefficient is $\sigma$=0.015 N/m, the oil phase viscosity is $\eta$ = 0.014 Pa.s, the tube radius of the micropipette tip is $R_{tube}$ = 75 μm, the density of the oil is 0.8 g/cm$^3$, the flow rate $Q$ of the aqueous phase can be adjusted from

approximately 10 nL/s to more than 1500 nL/s, the amplitude of the oscillation $A$ usually ranges from 0.4 mm to 0.6 mm, and the oscillation frequency is set to 120 Hz. At last, there is one parameter, the neck diameter $d_{neck}$, which is difficult to deal with. According to literature [3], $d_{neck}$ was usually found to be $d_{neck}$=0.4$R$. We use this result in the calculation as it is consistent with our experimental observation. As a result, we obtain that the *We* number, *We* ~ O(10), is almost two orders of magnitude larger than the *Ca* number, *Ca* ~ O(0.5). The large *We* number clearly manifests the inertial force-dominated mechanism, while the viscous effect is not sufficient to complete droplet segmentation as *Ca* < 1. More importantly, the *We* number in the OsciDrop system is even larger than the *Ca* number in the spinning/revolving system, indicating a stronger segmenting effect from the inertial force. We believe this is the key mechanism for our OsciDrop to generate highly uniform droplets.

**Text S5. Force analysis for the asymmetrical oscillation**

We establish an asymmetrical oscillation (see Fig. S2), connecting a long triangle wave (which lasts the first 4$T$/5 in each period) with a short cosinusoidal wave (which lasts the remaining $T$/5 in each period) in a period. We find that this asymmetrical oscillation is more efficient to achieve highly uniform droplet segmentation during each oscillation period. In such a configuration, the inertial force $F_i$ can be divided into two parts: when $0 < t < 4T/5$ (called the initial stage), $F_i = 0$ as the droplet displacement varies linearly with time $t$ and the acceleration equals to zero. When $4T/5 < t < T$ (called the segmenting stage), for a cosinusoidal wave with displacement $x(t) = -A\cos[\frac{5\pi}{T}(t-\frac{4}{5}T)]$, the acceleration $a(t)$ can be calculated by taking the derivative twice:

$$a(t) = A(5\pi/T)^2\cos[\frac{5\pi}{T}(t-\frac{4}{5}T)] \quad \text{[S8]}$$

Thus, the inertial force can be deduced as:

$$F_i = \rho\pi R_{tube}^2 u_f tA(5\pi/T)^2\cos[\frac{5\pi}{T}(t-\frac{4}{5}T)] \quad \text{[S9]}$$

As a result, we obtain Eq. 4 in the *Results* section to describe the inertial force. The viscous drag force and the interfacial tension are still described by Eq. (S3) and (S4) respectively.

We find that the triangle wave can serve as a "force switch" as its linear motion results in zero inertial force, and the cosinusoidal waves can be used to precisely determine the moment when the maximum inertial force should be applied. The reason is that, at the turning moment $t = 4T/5$ the inertial force suddenly reaches its maximum value according to Eq. S9. This maximum inertial force $F_i$ at the turning moment results in a sharp increase of the breaking force $F_i + F_v$ at the same moment. Therefore, we can precisely design when the segmenting occurs based on the turning moment connecting with a cosinusoidal wave. We will employ force plot explained in Text S5 below to highlight the turning moment and different working ranges. In addition, extending the length of the initial stage and reducing the length of the segmenting stage can enlarge the inertial force $F_i$.

**Text S6. Force plot and three working ranges**

Based on Eq. S3, S4 and S9, we calculate the temporal variation of the major forces in one oscillation period. Thus, a three-dimensional (3D) force plot among the interfacial tension $F_\sigma$, the viscous drag force $F_v$, and the inertial force $F_i$ can be provided to compare different effects. The force plots of the asymmetrical oscillation used in the current experiments have been shown in Fig. 2F. The $x$ and $y$ axes in the 3D plot are the elapsed time $t$ (ms) and the flow rate $Q$ (nL/s), and the vertical $z$ axis indicates the values of different forces. The interfacial tension $F_\sigma$ that resists droplet segmentation is displayed by the grey surface. According to Eq. S4, the interfacial tension $F_\sigma$ is constant and independent of $t$ and $Q$. The summation of the $F_i$ and $F_v$ that causes droplet break-off is shown by the colorful curved surface, in which red means larger force and blue means smaller force. The force comparison plot clearly shows that in one single period there is a region where $F_i + F_v > F_\sigma$. The corresponding time $t$ and flow rate $Q$ in such region indicate the condition to achieve a successful droplet segmentation. As mentioned above in Text S4, the sharp increase of the breaking force occurs just after the turning moment $t = 4T/5$, which is verified by the crossover of the colorful surface ($F_i + F_v$) above the grey horizontal plane $F_\sigma$.

The force plot helps understand the existence of three working ranges of the OsciDrop system. As indicated by the dashed lines in Fig. 2F, a lower threshold flow rate $Q_{th\text{-}L}$ can be clearly seen, below which $F_i + F_v$ is always smaller than $F_\sigma$ and the droplet segmentation does not occur in this period. When the oscillation frequency $f = 120$ Hz, the lower threshold flow rate is found to be about $Q_{th\text{-}L} = 20$ - 30 nL/s. If the frequency is reduced to 30 Hz, the $Q_{th\text{-}L}$ will increase to about 150 nL/s, implying an increasing difficulty to generate small droplet at low flow rate. Note that when flow rate $Q$ is large enough there is another threshold $Q_{th\text{-}U}$. Above this upper threshold $Q_{th\text{-}U}$, the breaking force ($F_i + F_v$) dominating droplet segmentation and the interfacial tension become comparable even in the initial stage, resulting in nonuniform droplet segmentation or generation of satellite droplets. When the oscillation frequency $f = 120$ Hz, the upper threshold flow rate is found to be about $Q_{th\text{-}U} = 1400$ nL/s. Thus, we conclude the monodisperse range (within the two dashed lines in Fig. 2F) in between the two threshold flow rates $Q_{th\text{-}L}$ and $Q_{th\text{-}U}$.

Based on these two threshold flow rates, we summarize a phase diagram of the droplet segmentation as shown in Fig. 2E. The three different working ranges are clearly displayed in the phase diagram. The perfectly monodisperse range is in the middle, whereas in the left one with smaller amplitude and flow rate the droplet segmentation frequency is usually half of the oscillation frequency, and in the right one with larger amplitude and flow rate the droplet is also nonuniform and the droplet segmentation frequency is larger than the oscillation frequency. We provide a blue dashed line and a red dashed line in the phase diagram (Fig. 2E) to theoretically predict the boundaries between different working ranges. This theoretical prediction is numerically calculated based on the corresponding crossover point between $F_i + F_v$ and $F_\sigma$. We can see good agreement between the prediction and the experimental observation (experimental results of different working ranges are categorized by different symbols in Fig. 2E).

**Text S7. Performance of sinusoidal oscillation and corresponding force plot**

We take sinusoidal oscillation as an example to show its force plot and compare it with asymmetrical oscillation. The displacement of a sinusoidal wave is $x(t) = A\sin(2\pi t/T)$, so the acceleration $a(t)$ can be calculated as:

$$a(t) = -A(2\pi/T)^2 \sin(2\pi t/T) \quad \text{[S10]}$$

And the inertial force is:

$$F_i = -\rho\pi R_{tube}^2 u_f t A(2\pi/T)^2 \sin(2\pi t/T) \quad \text{[S11]}$$

The negative signs in Eq. S10 and S11 only determine the direction. The equation of the inertial force $F_i$ under sinusoidal oscillation (Eq. S11) is similar to that under asymmetrical oscillation given in Eq. S9, implying that sinusoidal oscillation could also work with proper flow rate and oscillation frequency/amplitude. As shown in Fig. S6A, we can obtain highly uniform droplets from about 2 - 5 nL under a 120 Hz sinusoidal oscillation. To generate 1 nL or smaller droplets uniformly, we have to increase the frequency to 160 Hz (corresponding to decrease $T$ in Eq. S11) to enlarge the inertial force $F_i$ according to Eq. S11. Compared with Eq. S9, the $F_i$ of sinusoidal oscillation is smaller than that of the asymmetric oscillation, owing to the prefactor $(2\pi)^2$ in Eq. S11 instead of $(5\pi)^2$ in Eq. S9. We show the force plot under 120 Hz sinusoidal oscillation using low flow rate in the left of Fig. S6B. We can see that the lower threshold flow rate $Q_{th\text{-}L}$ is about 180 nL/s, below which the breaking force is not large enough to overcome interfacial tension. According to Eq. 3, this lower threshold $Q_{th\text{-}L}$ means that the smallest droplet that can be obtained under 120 Hz oscillation is about 180/120 = 1.5 nL. If we further increase the flow rate to be more than 1000 nL/s, we can see from Fig. S6A that droplet segmentation is achieved twice in one oscillation period. To explain this phenomenon, we calculate the temporal variation of the breaking force. In the calculation, if the droplet segmentation complete when the breaking force exceeds the interfacial tension, the elapsed time $t$ in Eq. S11 will reset to zero to simulate the start of another droplet segmentation. The force plot for large flow rate is shown in the right of Fig. S6B, in which we can see two similar peaks of the breaking force in both half of the oscillation period. The force plot predicts that when the flow rate is larger than approximately 1200 nL/s, we will obtain double droplet segmentation in one oscillation period. The prediction is consistent with the experimental observation shown in Fig. S6A.

**Text S8. Working principle and droplet generation cycle using the standard 10-μL micropipette tip**

In addition to generate monodisperse droplets using customized micropipette tips with an i.d. 100~150 μm, we also successfully generate monodisperse nanoliter droplets using the standard 10 μL micropipette tip with an i.d. 400 μm. Due to the relatively large inner diameter of the tip orifice, the interfacial tension is dominant at the beginning (called pre-initial stage). This pre-initial stage lasts a certain time until the aqueous phase protrudes into the oil phase and the interfacial meniscus reaches equilibrium dominated by the interfacial tension. After the pre-initial stage, the droplet segmentation process becomes similar to that using customized micropipette tips mentioned before, which contains two stages in a repeatable period: During a long initial stage, the aqueous stream bulges and elongates in the oil phase, and is segmented into monodisperse droplets during a short segmenting stage. The droplet generation process is schematically illustrated in Fig. S3. The droplet segmentation is done with the assistance of the sharp edge of the orifice wall rather than tip dimensions. Successful generation of monodisperse nanoliter droplets using standard 10 μL micropipette tip provides a good example to demonstrate the robustness of the OsciDrop system.

**Text S9. Investigation of the effect of wall thickness on the droplet generation using an one-side tapered tip with OsciDrop**

The idea that the droplet segmentation is strongly influenced by edge of the orifice wall inspires us to investigate the effect of the wall thickness. In order to clarify the effect of the wall thickness on the droplet segmentation, we fabricate a one-side tapered micropipette tip by cutting part of the wall carefully with the aid of microscopic observation. The tapered tip is shown in Fig. S5A. We suppose that the droplet segmentation would occurs in the thin-wall side, as schematically illustrated in Fig. S5B. The reasons can be understood as follows: (1) The sharp edge of the thin wall faciliate segmenting the aqueous stream. (2) According to Eq. S9, the inertial force scales as $F_i \sim \rho Q t = \rho V(t)$, where $V$ is droplet volume. Once the droplet segmentation due to the sharp edge effect occurs on the thin-wall side, after half period on the thick-wall side the droplet hasn't grown to a sufficiently large volume, causing the inertial force $F_i$ on the thick-wall side is usually smaller than that on the thin-wall side. We perform experiments using one-side tapered tip at flow rates of 120 nL/s and 600 nL/s under oscillation frequency $f$ = 120 Hz. The results verify our assumption that droplet segmentation occurs on the thin-wall side (Fig. S5C) rather than on the thick-wall side (Fig. S5D). The droplet segmentation can greatly benefit from the sharp edge effect, which does not require a high oscillation speed to assist segmenting. Thus, this strategy will be very helpful to reduce the possibility of generating satellite droplet. In particular, we find that this one-side tapered tip works extremely well for generating picoliter droplets (Fig. S5E). We successfully obtain very uniform droplets with volume ranging from 200 - 800 pL using such tapered tip, without generating noticeable satellite droplets.

### Supporting Tables

**Table S1.** Primers for African swine fever virus (ASFV) detection with dLAMP

| Primer | Oligonucleotide sequence |
|---|---|
| FIP | 5'-GCCTCGTTGGTGGAAAGGATCCTTCAACGCATAGGAGAC-3' |
| BIP | 5'-TGGCGGGAGAGGAAGGAACTATGACGGCGTAATGAAGAT-3' |
| F3 | 5'-GCATGTACTACAACGATTAGGA-3' |
| B3 | 5'-AAGTCGCCGATTTGGTTT-3' |

**Table S2.** Three primary control parameters ($Q$, $A$, $f$) for generating monodisperse droplets spanning picoliter to microliter range

| Expected droplet volume $V$ | Flow rate $Q$ | Oscillating frequency $f$ | Oscillating amplitude $A$ |
|---|---|---|---|
| 200 pL | 24 nL/s | 120 Hz | 0.77 mm |
| 500 pL | 60 nL/s | 120 Hz | 0.75 mm |
| 1 nL | 120 nL/s | 120 Hz | 0.55 mm |
| 5 nL | 500 nL/s | 100 Hz | 0.54 mm |
| 10 nL | 1000 nL/s | 100 Hz | 0.47 mm |
| 50 nL | 1000 nL/s | 20 Hz | 0.75 mm |
| 100 nL | 500 nL/s | 5 Hz | 1.14 mm |
| 500 nL | 500 nL/s | 1 Hz | 1.14 mm |
| 1 μL | 500 nL/s | 0.5 Hz | 1.14 mm |
| 2 μL | 1000 nL/s | 0.5 Hz | 1.14 mm |

**Supporting Figures:**

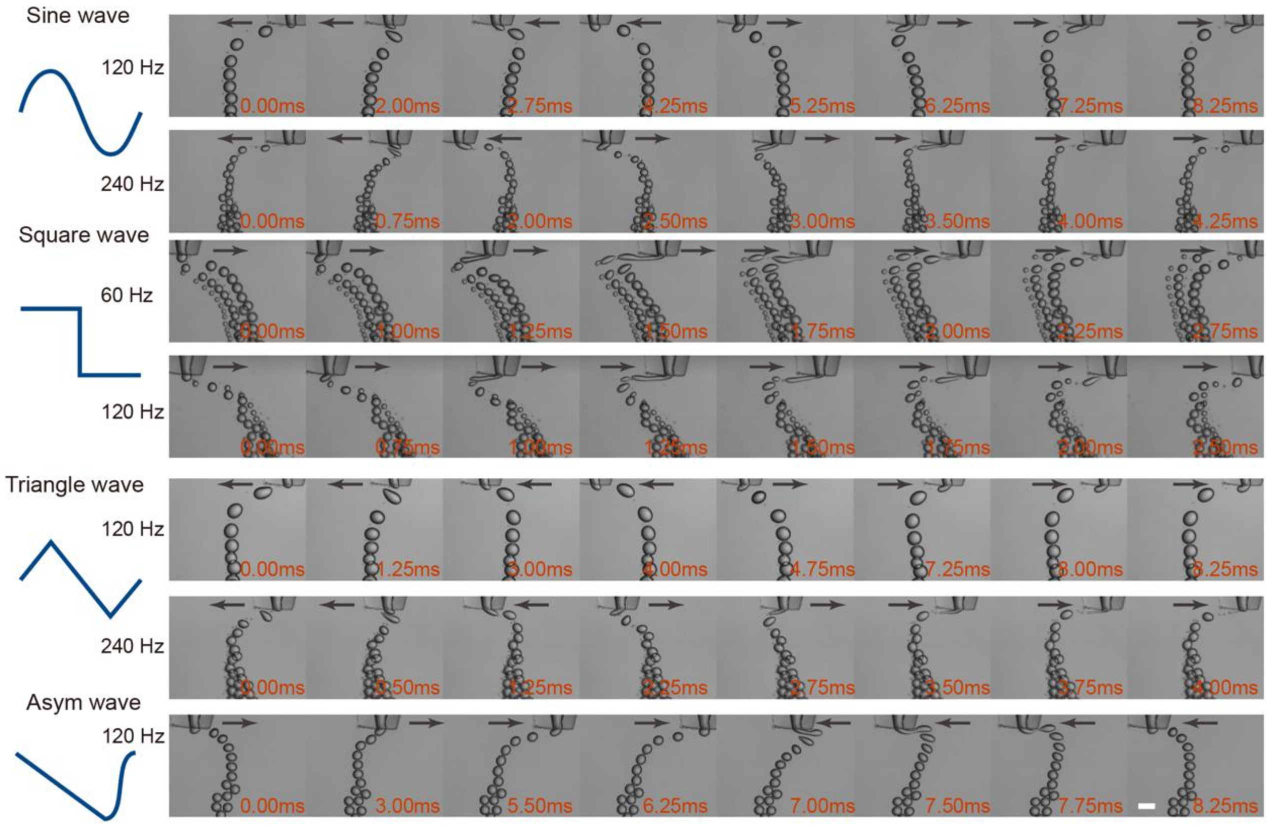


**Fig. S1** Droplet generation using different oscillation waveforms, including sine, square, triangle, and asym wave, under different oscillation conditions (only part of results were shown herein). Scale bar is 200 µm.

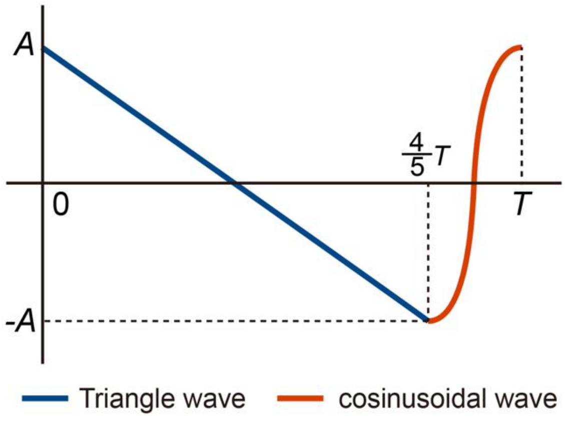


**Fig. S2** The asymmetrical oscillation waveform consists of a long triangle wave (called initial stage that lasts the first 4*T*/5 in each period *T*) and a short cosinusoidal wave (called segmenting stage that lasts the remaining *T*/5 in each period *T*).

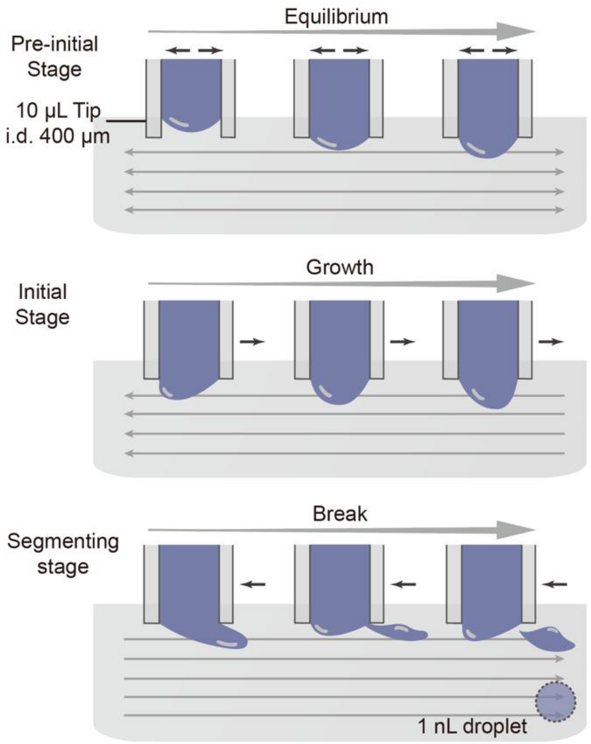


**Fig. S3** A schematic illustration of the working principle and a droplet generation cycle using the standard 10 µL micropipette tip. The droplet generation process includes a pre-initial stage that the head of the aqueous stream cannot be segmented into droplets as interfacial tension is dominant. After pre-initial stage, the droplet segmentation begins. During a long initial stage, the aqueous stream bulges and elongates in the oil phase, and is segmented into monodisperse droplets during a short segmenting stage. Black arrows indicate the moving direction of the micropipette tips. Grey arrows represents the direction of relative motion of the oil phase.

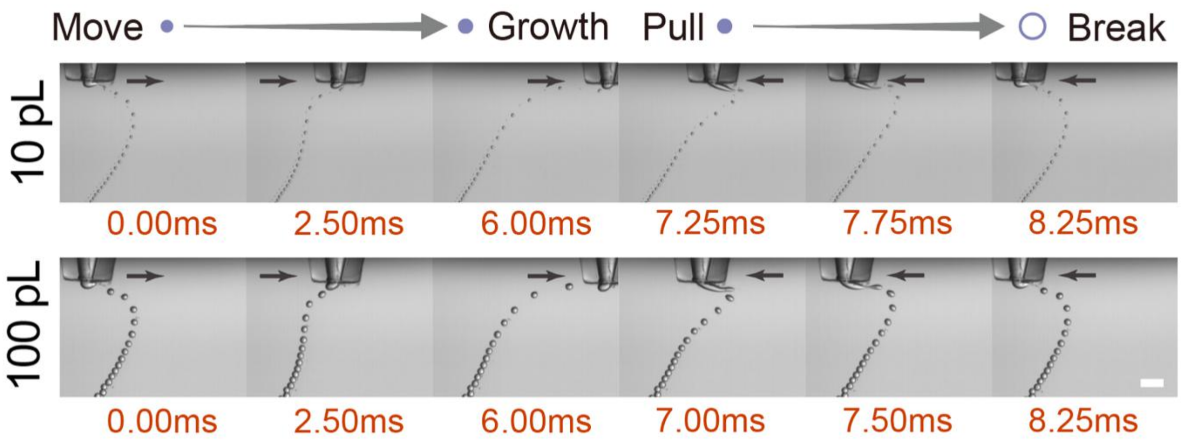


**Fig. S4** Time-series microscopic images of the droplet generation cycle (expected volumes of 10 pL and 100 pL) under 120 Hz asymmetrical oscillation. Black arrows indicate the moving direction of the micropipette tip. Scale bar is 200 μm.

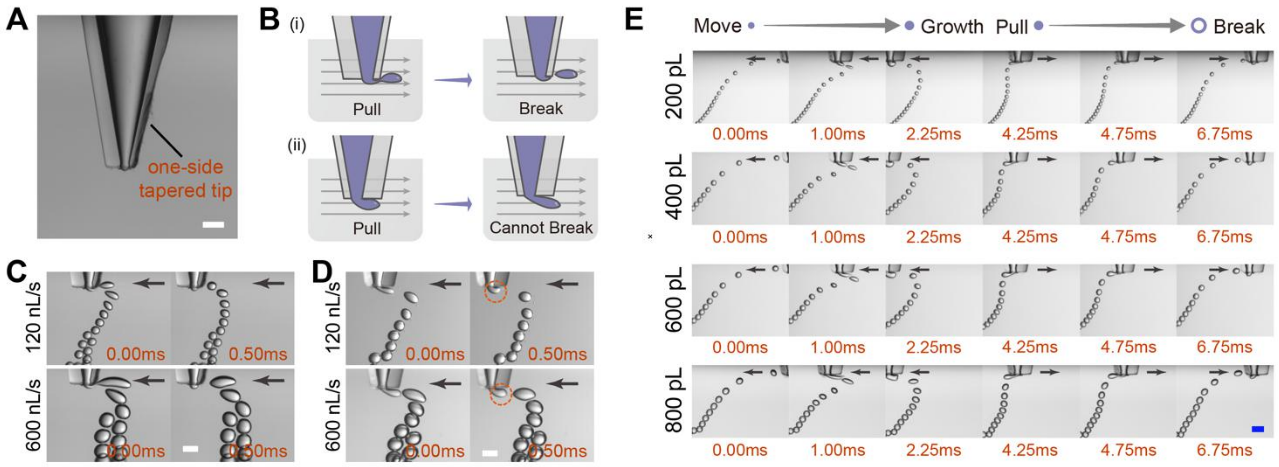


**Fig. S5** Droplet generation using a one-side tapered tip with OsciDrop. (A) Microscopic image of the one-side tapered tip. (B) Schematic illustrations of the feasibilty of segmenting droplets using different sides of the one-side tapered tip. Droplet segmentation at flow rates of 120 nL/s and 600 nL/s can (C) /cannot (D) be achieved using different sides of the one-side tapered tip. (E) Facilitating picoliter droplet generation using one-side tapered tip. Black arrows indicate the moving direction of the micropipette tip under the air/oil interface. Scale bar is 200 μm.

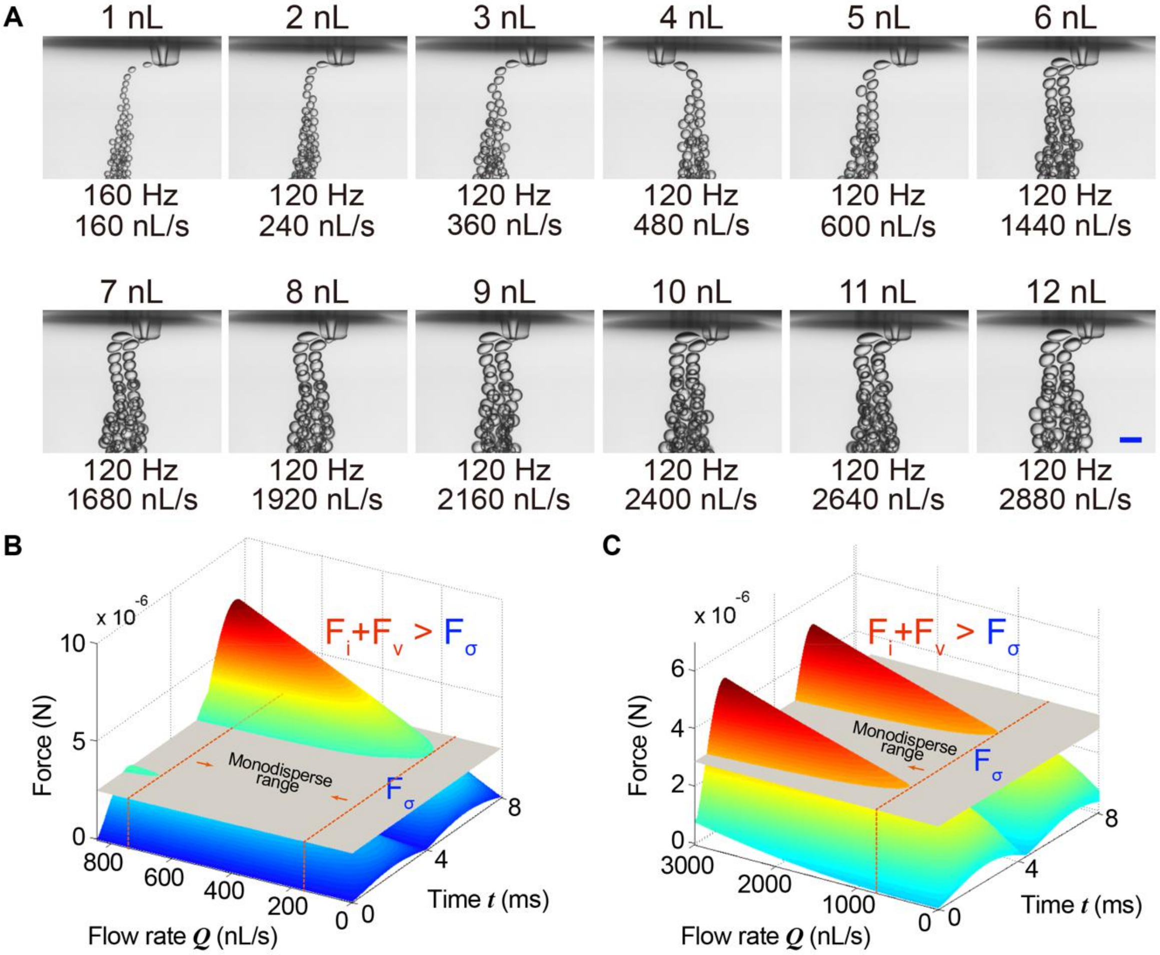


**Fig. S6** Droplet generation under sinusoidal oscillation with OsciDrop. (A) Size-tunable generating monodisperse nanoliter droplets (1 nL to 12 nL) under sinusoidal oscillation using OsciDrop. Scale bar is 400 μm. (B) Theoretical force comparison three-dimensional (3D) maps showing the temporal variations of $F_i + F_v$ (colorful curved surface) and $F_\sigma$ (grey surface) during one period $T$ ($f$ = 120 Hz) at low flow rates (B *left*) and high flow rates (B *right*).
